# No effect of anodal high-definition transcranial direct current stimulation during motor sequence learning in older people

**DOI:** 10.64898/2026.02.18.706569

**Authors:** Silke Kerstens, Sanne Broeder, Moran Gilat, Evelien Nackaerts, Britt Vandendoorent, Alice Nieuwboer, Jean-Jacques Orban de Xivry

**Author notes:** **Corresponding author**, Jean-Jacques Orban de Xivry.

## Abstract

**Background:** Transcranial direct current stimulation (tDCS) targeting the primary motor cortex (M1) has previously been shown to enhance motor learning in older adults. However, findings across studies are inconsistent, which may be partly due to variability in stimulation parameters and montages across experimental protocols.

**Objective:** We aimed to replicate the effect of 1mA conventional anodal tDCS over the M1 hotspot during one session of motor sequence learning on retention (24h) compared to sham stimulation in older adults. In addition, we aimed to compare the effect of high definition (HD tDCS) versus conventional tDCS on motor sequence learning to explore if stimulation montage and according focality affects the effect sizes found.

**Methods:** In a pre-registered, double-blind, randomized, sham-controlled, parallel study design including 52 older adults, we investigated the effects of conventional and HD tDCS on motor sequence learning in a serial reaction time task (SRTT), using a crossover design for the stimulation montage.

**Results:** Although all groups showed motor sequence learning over time, both during practice sessions (main effect of time: p < 0.0001), as well as across session (main effect of time: p < 0.0001), we observed no significant effects on learning between stimulation groups (main effect of stimulation: p = 0. 68) or montages (main effect of montage: p = 0. 66). For all groups, motor sequence learning improvements were maintained but not further enhanced after 24 hours of consolidation (HD tDCS: p = 0.64; conventional tDCS: p = 0.76; HD sham: p = 0.69; conventional sham: p = 0.57).

**Conclusion:** Our findings indicate that neither conventional nor HD tDCS enhanced motor sequence learning in older adults. To better understand potential long-term or cumulative effects, we recommend that future studies investigate the effects of repeated tDCS sessions administered throughout the motor learning process.

## Introduction

Neuroplasticity refers to the ability of the central nervous system (CNS) to modify its structure and function in response to stimuli (Pascual-Leone et al. 2005), and plays an essential role in the capacity for learning. However, with ageing neuroplasticity gradually declines (Oberman and Pascual-Leone 2013). Previous work has shown that sequential motor learning, for example, which involves the improvement of spatial or temporal accuracy of motor sequences with practice (Dahms et al. 2020), is impaired in older adults (King et al. 2013). After skill acquisition, the sequential motor skill is encoded in specific brain areas (Pascual-Leone et al. 1995), such as in the primary motor cortex (M1), amongst others, (Pascual-Leone et al. 2005) during a process referred to as consolidation (Kantak and Winstein 2012). This process involves changes of neural excitability and plasticity in these regions and if consolidation is successful, the improvements are maintained over time. In other words, the learned skill is maintained (Kantak and Winstein 2012). As consolidation of motor sequence learning involves greater integration within a cortico-striatal functional network (Debas et al. 2014), impairments in this network would imply a decreased ability for consolidation. Therefore, age-related alterations in brain connectivity within motor learning networks may explain the reduced consolidation abilities observed in older adults (King et al. 2013).

To counteract this natural decline or to optimize rehabilitation during recovery from an injury (Kleynen et al. 2020), researchers have been exploring methods to improve neuroplasticity in older adults. In this context, transcranial direct current stimulation (tDCS) has been proposed as a method to potentially boost motor learning (Buch et al. 2017; Orban de Xivry and Shadmehr 2014), albeit with inconsistent results (Jalali, Chris Miall, and Galea 2017; Kaminski et al. 2024; Vandendoorent et al. 2023). Transcranial direct current stimulation (tDCS) is a noninvasive brain stimulation technique that involves the application of a small direct current of 1-4 mA via stimulation electrodes on the scalp (Nitsche et al. 2000; Nitsche and Bikson 2017), with the positively charged electrodes referred to as anodal and the negatively charged electrodes as cathodal. These electrodes are strategically positioned above brain regions that play a role in motor learning processes such as the M1, among others (DaSilva et al. 2011). When anodal electrodes are positioned over the target region, they deliver a positive charge, a process known as anodal tDCS.

In conventional tDCS, stimulation is typically applied by two large conductive rubber electrodes. Rather than being placed directly on the scalp, these electrodes are usually embedded in saline-soaked sponges to enhance conductivity and minimize skin irritation during stimulation. However, previous studies have indicated that this method stimulates a broad area underneath the stimulation electrodes (Caparelli-Daquer et al. 2012) and some propose that most of the applied current is shunted transcutaneously before reaching the targeted cortical regions (Khatoun et al. 2018).

In an attempt to enhance focality, a new stimulation method has been developed in which a smaller stimulation electrode is placed over the targeted area and four reference electrodes are arranged in a circle around this area, referred to as high definition tDCS, or HD tDCS (Bortoletto et al. 2016; Villamar, Volz, et al. 2013). Based on computational modeling studies, it has been projected that this montage results in a higher electric field in the targeted cortical regions directly underneath the stimulation electrodes (Villamar, Wivatvongvana, et al. 2013), thereby inducing more focalized stimulation. However, few studies directly compared HD tDCS with conventional applications during motor learning (Firouzi et al. 2023a) and if so, mostly in young healthy adults.

Furthermore, the effects of tDCS, regardless of application mode, can be attributed to both transcranial and transcutaneous mechanisms (van Boekholdt et al. 2021). The transcranial mechanism refers to the depolarization of membrane potentials of cortical neurons during tDCS (van Boekholdt et al. 2021), whereas the transcutaneous mechanism suggested that indirect stimulation of brain circuits occurs via the stimulation of peripheral nerves in the scalp (van Boekholdt et al. 2021).

In a previous study, we found that applying conventional tDCS during writing practice in patients with Parkinson’s disease (PD) led to greater improvements in skill level compared to sham stimulation (Broeder et al. 2023). Although our recent meta-analysis in older adults with and without PD revealed that the additional effects of tDCS were variable and highly task-dependent (Vandendoorent et al. 2023), we hypothesize that tDCS applied during motor sequence learning in healthy older adults may also support better retention of motor sequence learning over time. To test this hypothesis, the present study investigated the effect of conventional anodal tDCS over M1 on motor sequence learning and retention 24h after a single practice session compared to sham in older adults in a parallel design. In addition, we also investigated whether an extra session of continued learning changed the outcomes. Finally, we explored the surplus effects of stimulation montage by comparing HD tDCS to conventional tDCS using an additional crossover condition within the overall parallel design.

## Methods

In a double-blind, randomized, sham controlled, parallel study design including 52 older adults, we investigated the effect of conventional and HD tDCS on motor sequence learning in a serial reaction time task (SRTT), with an additional crossover design for stimulation montage (Figure 1A).

**Figure 1.**
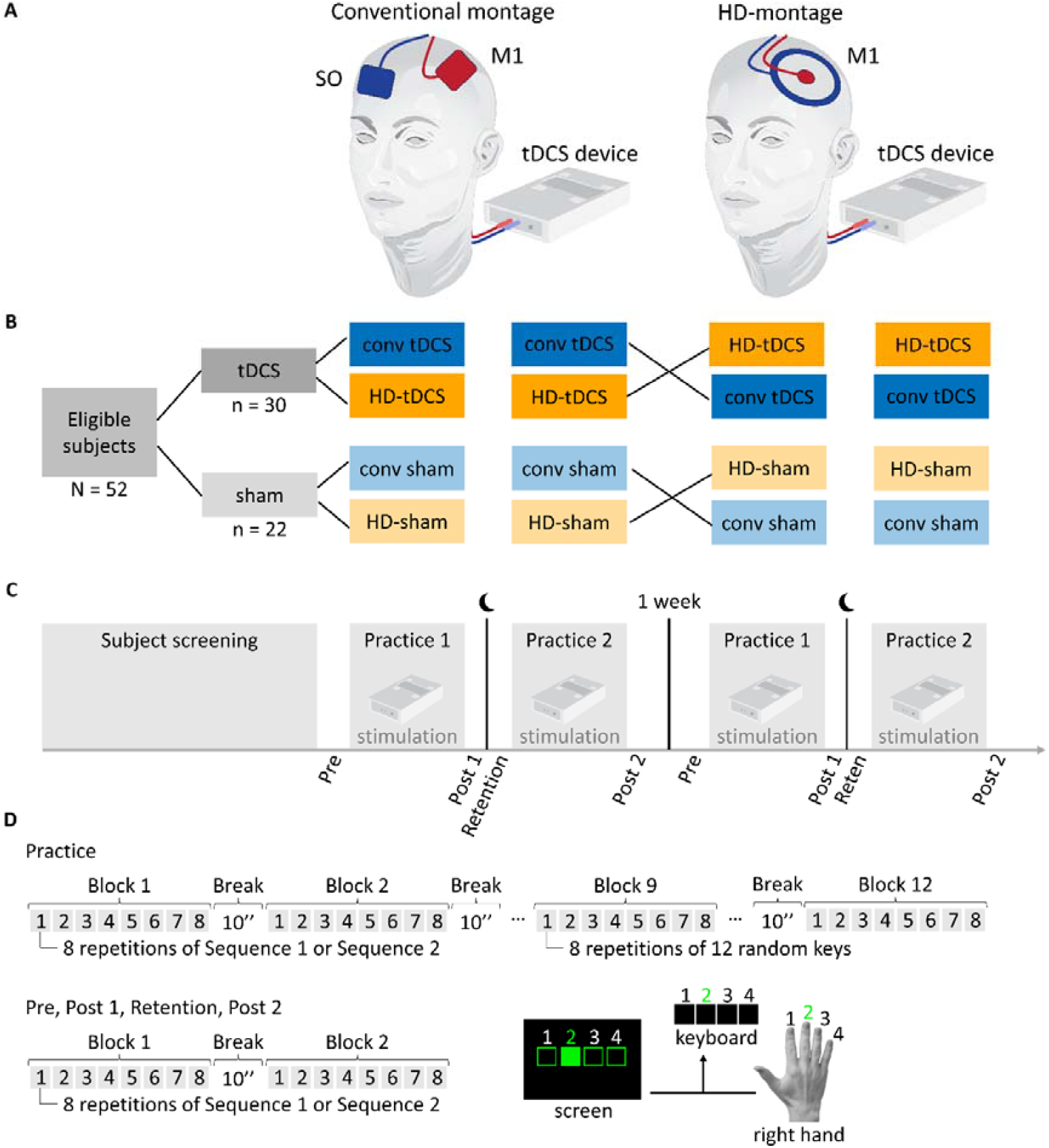
Study protocol. (A) Conventional and HD tDCS montage. (B) A total of 52 older adults participated in the study, with 30 randomly assigned to the tDCS group and 22 to the sham group. Regardless of stimulation group, all subjects experienced both montages in a crossover design, with the order of montages randomized. (C) After screening, all subjects completed a four-session motor sequence learning paradigm consisting of two phases, with each phase including two sessions conducted on consecutive days. One week interval between the two phases was implemented to prevent carryover effects. Each phase consisted of a two-session SRTT motor sequence learning paradigm. Subjects first established a baseline (Pre), practiced the SRTT sequence with concurrent stimulation, and were tested (Post 1) afterwards without stimulation. After a 24-hour consolidation period, retention was assessed (Retention) at the beginning of the second session. Next, they continued practicing the same sequence with stimulation and were tested again without (Post 2). (D) The SRTT is a motor sequence learning task in which the four squares on the screen correspond to four keys on the keyboard, and to the four fingers of the right hand. Subjects were instructed to respond as fast and as accurately as possible to the filled squares by pressing the corresponding key on the keyboard. Each subject practiced two 12-key sequences across the two phases of the paradigm, with the sequence order randomly assigned. Each practice session included 88 repetitions grouped into 11 blocks of 8 repetitions, requiring 96 key presses per block. To prevent fatigue, a 20-second break was provided between blocks, indicated by red squares on the screen.

### Ethical approval

This study was approved by the Ethics Committee Research UZ/KU Leuven (S60893) and pre-registered on <u>Open Science Framework</u>. All subjects received financial compensation of €50 for their participation and their travel expenses were reimbursed. Data collection occurred between July 2020 and March 2022.

### Subjects

In total, 52 older adults of on average 68.4 ± 8.0 years old (M: 27 and F: 25) were included based on an a priori sample size calculation (see below) and according to the following inclusion criteria: age ≥ 50 years old, right-handedness determined by the Edinburgh Handedness Inventory (EHI), cognitively-intact based on the Mini Mental State Examination (MMSE) score ≥ 24, and 4) no motor- or neurological disorders or implants. After inclusion in the study, subjects were randomly allocated to either the stimulation group (tDCS) or the sham group. After study inclusion, an independent researcher carried out a concealed randomization procedure to assign the subjects to either the active tDCS or the sham stimulation group using an online randomization tool (<u>Simple randomization service</u>, Sealed Envelope, 2017). Regardless of the stimulation group, each subject also underwent two montages: the conventional and the HD tDCS montage, with the order randomly assigned (Figure 1B).

### Experimental protocol

Prior to inclusion, all subjects underwent screening during an initial session. If the subjects met all the initial eligibility criteria, they proceeded to the clinical tests to assess the additional inclusion criteria before proceeding to the next phase. Those who met all inclusion criteria were included in the study and provided with a detailed explanation of the protocol, before signing the informed consent form. Afterwards, subjects were familiarized with the SRTT task and then randomly assigned to either the active tDCS group (n_tDCS_ = 30) or the sham group (n_sham_ = 22) using permuted blocks of four, stratified by age (<65 vs. ≥65 years) and gender.

Regardless of stimulation group, all subjects completed a motor sequence learning paradigm consisting of two phases (Figure 1C), with a one-week washout period between the phases. Within each phase, subjects performed a two-session SRTT motor sequence learning paradigm conducted on consecutive days. At the start of each phase, baseline performance was established (Pre). During the subsequent practice session, subjects practiced a motor sequence while receiving either tDCS or sham stimulation. At the end of the session, their improvements in motor sequence learning were assessed in a test (Post 1) with no stimulation applied. Following a consolidation period, retention was assessed through a retention test (Retention) 24 hours later. Subsequently, subjects continued to practice the same motor sequence in a second session and were then assessed in a final test (Post 2) to examine ‘continued learning’ (Figure 1C).

After a one-week washout period, the same protocol was repeated using the alternate electrode montage. In the first phase, subjects were randomly assigned to receive either the conventional or the HD montage, following the CONSORT guidelines (Schulz, Altman, and Moher 2010). During the second phase of the experiment, the alternate montage was applied. The one-week interval between the two phases was included to minimize carryover effects.

### tDCS and sham stimulation

During active tDCS, a small direct current of 1mA was administered using the DC-Stimulator (NeuroConn GmbH, Germany) over the left primary motor cortex (M1) for 20 minutes during practice at the optimal scalp position (hotspot). The hotspot was determined individually for each subject by applying single-pulse transcranial magnetic stimulation (TMS) using the Magstim (BiStim2, UK, figure-of-eight coil, 70 mm diameter) to elicit a motor response in the right first dorsal interosseous (FDI) muscle, as measured by surface electromyography (Bagnoli-4, Delsys Inc., USA) (Rossini et al. 2015). This location was standardized across sessions using a head mold, as used in previous work (Broeder et al. 2019). The subjects and the experimenter were blinded to the type of stimulation (anodal or sham) through the ‘study mode’ of the tDCS stimulator (DC-Stimulator, NeuroConn GmbH, Germany). In this mode, the blinded researcher used a 5-digit number provided by an independent researcher, which encoded the type of stimulation. The active current was gradually increased to 1 mA over 10 seconds, and after the practice session, it gradually decreased to zero. The sham current was ramped up to 1 mA over 10 seconds and held constant for another 10 seconds to mimic the sensation of active stimulation, after which it was ramped down to zero over an additional 10 seconds. In each session, impedance was verified to be below 20 kΩ prior to initiating stimulation and continuously monitored throughout the stimulation period to ensure reliable stimulation and safety. Stimulation was administered by the same blinded researcher in all sessions.

### Conventional and HD tDCS

Conventional tDCS was applied over two rectangular electrodes of 5×7 cm with a current density of 0.03 mA/cm^2^. The anodal electrode was placed over the left M1 hotspot, and the cathode electrode over the right supraorbital (SO) region (Figure 1A). In HD tDCS, the central anodal electrode was applied over the left M1 hotspot using a circular electrode with a diameter of 2.5 cm and a current density of 0.2 mA/cm^2^, surrounded by a ring-shaped cathode electrode with a distance of 3.25cm between the central anodal and the circular cathodal electrode (Figure 1A). The researcher was not blinded for montage.

### Blinding

Immediately after stimulation, subjects were asked to rate the stimulation sensation on a visual analogue scale (VAS) from 0 to 10, with higher scores representing a higher intensity. Additionally, at the end of the last practice session, both the subjects and the researcher were asked to guess whether the subject had received active tDCS or sham stimulation throughout the experiment.

### Serial reaction time task

The SRTT is a motor sequence learning task in which subjects improve their ability to execute a sequential order of motor actions. Subjects are asked to respond to visual stimuli presented on a screen by pressing a corresponding key on a keyboard. In this study, subjects learned to press four keys in a sequential order using the four fingers of the right hand, excluding the thumb. The four fingers corresponded to four empty squares on the screen, and to four keys on the keyboard (Figure 1D). During the task, the visual stimulus was presented by filling one of the squares. Upon presentation of the stimulus, the subjects were instructed to respond as fast and as accurately as possible by pressing the corresponding key with the corresponding finger. Upon a key press, irrespective of correctness, the next stimulus was immediately presented on the screen. To assess performance during the task, we measured both response time (primary outcome), i.e. the time between the representation of the stimulus on the screen and the key press, and accuracy. To facilitate learning, all subjects were made aware of the presence of a sequential order in the task. However, no information was shared about the length of the sequence, or the number of repetitions.

Each subject practiced the following two 12-key sequences: Sequence 1) 4-2-3-4-3-2-1-4-1-2-3-1, or Sequence 2) 2-4-1-4-2-1-3-2-4-3-1-4 across the two phases of the paradigm respectively, with the order of the sequences randomly assigned. In each practice session, the designated sequence was repeated 88 times. However, to prevent fatigue, the repetitions were grouped into 11 blocks of 8 repetitions, with in total 96 key presses per block. Between blocks, a 20-second break was provided, during which the squares on the screen turned red. To determine a potential carryover effect from the first to the second phase of the paradigm, and to rule out the possibility that improvements over time are due to general skill enhancement rather than sequence learning, a random block was inserted between the 8th and 9th blocks in each practice session. In this random block, 96 key presses were presented in a pseudo-randomized order.

Performance tests were administered at the beginning and at the end of each practice session to monitor improvements over time. Each test consisted of two blocks of 8 repetitions of the sequence learned in that particular phase. The first test was used to establish the subjects’ baseline performance (Pre). Immediately after the first practice session, and a 2-minute break, the subjects’ improvements from that session were assessed through a second test (Post 1). Retention of practice, following a 24-hour consolidation period, was assessed through a third test (Retention). This was performed at the beginning of, and prior to, the second practice session. Analogous to the first session, continued learning during the second practice session was evaluated by a fourth test (Post 2) at the end of the session, following a 2-minute break.

## Data analysis

All data analyses were performed in Matlab R2024a (MathWorks, Natick, MA, USA), including the preprocessing, data analysis, and statistical analysis at a significance level of α < 0.05. Data analysis was carried out without knowledge of group allocation, as unblinding occurred only after the analysis.

### Sample size estimation

We estimated the sample size based on earlier study (Gbadeyan et al. 2016; Rumpf et al. 2017) with a calculated effect size of f = 0.12, derived from the effect of active anodal tDCS of M1 compared to sham combined with a finger sequence learning task in healthy older adults. Using G*Power software, a target sample of 24 subjects per group was determined to reach a power of 80% with an alpha of 5% in a general linear model design. We anticipated a drop-out rate of 10%. Therefore, a total of 26 subjects per group were included.

### Serial reaction time task

Motor sequence learning in the SRTT was analyzed based on linear mixed models in terms of mean reaction times per block, including stimulation (tDCS or sham), montage (conventional or HD), and time as fixed effects, with montage as a within-subject factor and stimulation as a between-subject factor. We decided to focus on mean reaction time as the primary outcome measure. Accuracy did not provide additional insight for the interpretation of the findings. To account for individual differences in baseline performance, we included subject as a random effect and likewise included the SRTT sequence as a random effect.

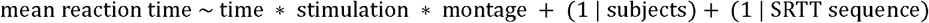

#### Effect of stimulation and montage on learning and retention

For each linear mixed model, we first calculated a mean reaction time for each timepoint Pre, Post 1 and Retention over the two blocks per test for each subject, and investigated the effect of stimulation, montage, and time on the mean reaction times per test. In case of significant effects, we analyzed if the observed effects were consistent over time and between groups in post-hoc tests. We first tested the groups for normality based on the one-sample Kolmogorov-Smirnov test. Accordingly, we compared the different timepoints using a two-sample t-test in case the samples were normally distributed. In case of abnormal distributions, we employed the Wilcoxon Signed-Rank test and the Mann-Whitney U test (in case of independent significantly different distributions).

#### Continued learning in the second practice session

To investigate continued learning in the second practice session, we analyzed the offline effect of stimulation and montage over time, comparing Retention and Post 2. For this linear mixed model, we first calculated a mean reaction time for each timepoint over the two blocks per test for each subject, and investigated the effect of stimulation, montage, and time on the mean reaction times per test, as above.

#### Online improvements during stimulation

As outlined in the preregistration, we explored the online effects of stimulation and montage on motor sequence learning over time during practice and concurrent stimulation. For this linear mixed model, we included the mean reaction times of all subjects including all blocks over the course of the first and second practice sessions, except the random block 9 in each session.

#### Carryover effect

To ensure that the outcomes in the second phase of the experiment were not confounded by residual learning effects from earlier practice sessions in the first phase of the experiment, a washout period of one week was implemented. To verify the effectiveness of this washout, we assessed baseline performance across phases by comparing mean reaction times in the first two sequenced blocks (Pre) of each initial practice session before and after the crossover in a linear mixed model with stimulation, montage and time as fixed factors, and subject and SRTT sequence as a random factor.

#### Blinding

We analyzed the effect of subject blinding on the reported stimulation intensities on the VAS over time based on a linear mixed model including stimulation (tDCS or sham), montage (conventional or HD), and time as fixed effects, montage as a within-subject factor, and stimulation as a between-subject factor. For this analysis, we counted the number of subjects who correctly identified their stimulation condition and calculated the success percentages to compare them between stimulation groups.

## Results

A total of 52 participants were included in the study and randomized to either the active tDCS group (n_tDCS_ = 30) or the sham group (n_sham_ = 22). The groups were comparable in terms of age (active tDCS = 69.03 ± 7.49 years, sham = 67.55 ± 8.82 years), gender distribution F/M (active tDCS: 15/15, sham: 10/12), and cognitive function (MoCA scores: active tDCS = 27.03 ± 2.03; sham = 26.73 ± 2.14). Detailed participant demographics and characteristics are provided in the Supplementary Materials in Table S1.

### Effect of stimulation and montage on learning and retention

The linear mixed model and figure 2 revealed a significant main effect of time (main effect of time: β = −44 ms, SE = 8.69 ms, t(298) = −5.10, p < 0.0001, 95% CI [−61 ms, −27 ms]), indicating that participants exhibited motor sequence learning. Performance improved from baseline to Post 1 and was maintained at Retention across all groups, as reflected in mean reaction times for conventional tDCS (611 ± 146 ms, 509 ± 155 ms, 520 ± 135 ms), conventional sham (588 ± 186 ms, 475 ± 148 ms, 512 ± 160 ms), HD tDCS (625 ± 167 ms, 559 ± 157 ms, 564 ± 148 ms), and HD sham (591 ± 145 ms, 514 ± 114 ms, 511 ± 107 ms), respectively (Figure 2). However, no significant effects were observed for stimulation (main effect of stimulation: β = 19.1 ms, SE = 45.7 ms, t(298) = 0.42, p = 0.68, 95% CI [−71.0 ms, 109.1 ms]) or montage (main effect of montage: β = −11.6 ms, SE = 26.1 ms, t(298) = −0.45, p = 0.66, 95% CI [−63.0 ms, 39.7 ms]). Subsequent post-hoc testing revealed that all groups showed significant improvements immediately after practice (HD tDCS: p = 0.013, conventional tDCS: p = 0.005, and HD sham: p = 0.047) compared to baseline, except for the conventional sham group (p = 0.074), which showed changes that did not reach statistical significance.

**Figure 2.**
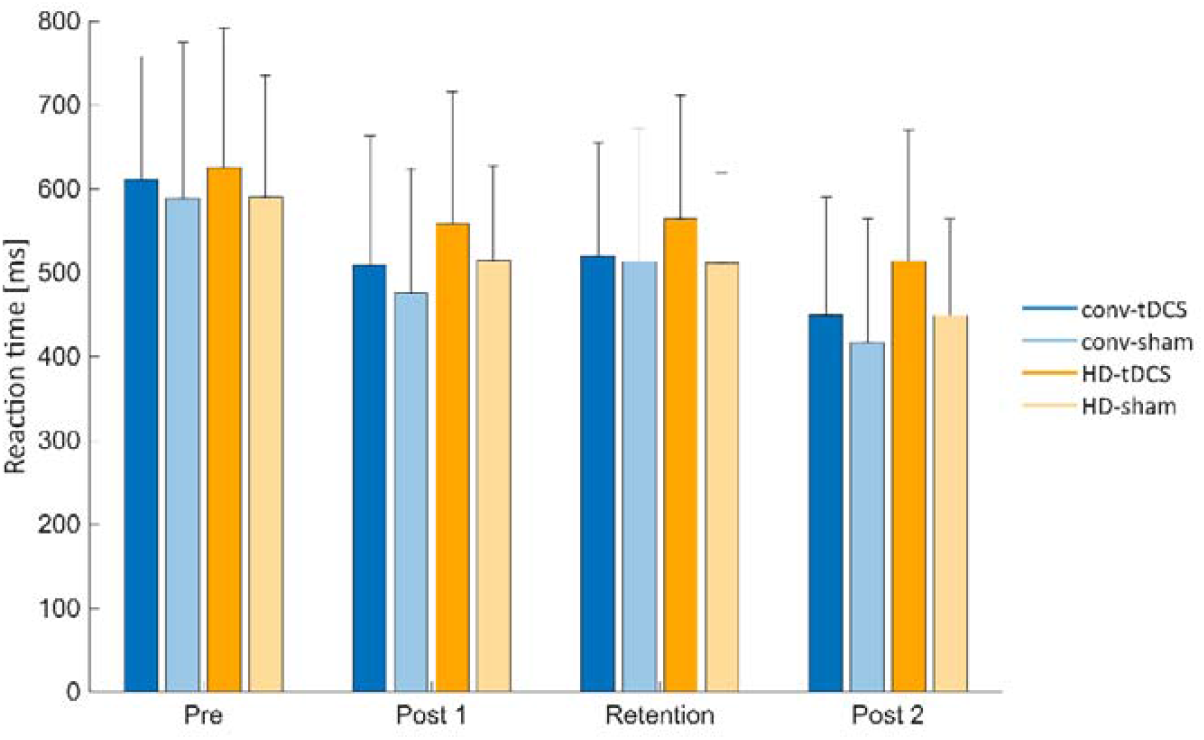
Motor sequence learning across sessions. The mean reaction times in milliseconds [ms] at the baseline (Pre), immediately after the first practice session (Post 1), after 24h-consolidation (Retention), and immediately after the second practice session (Post 2) for conventional tDCS (dark blue), conventional sham (light blue), HD tDCS (orange), and HD sham (yellow). The error bars indicate the standard deviation from the mean reaction times at group level.

When comparing performance at Retention to immediately after practice no significant differences were observed (HD tDCS: p = 0.64; conventional tDCS: p = 0.76; HD sham: p = 0.69; conventional sham: p = 0.57), suggesting that learning effects were maintained but not further enhanced after 24 hours without practice (Figure 2). This was also reflected in the mean reaction times for conventional tDCS (509 ± 155 ms versus 520 ± 135 ms), conventional sham (475 ± 148 ms versus 512 ± 160 ms), HD tDCS (559 ± 157 ms versus 564 ± 148 ms), and HD sham (514 ± 114 ms versus 511 ± 107 ms), respectively.

Although improvements were observed over time (main effect of time: β = −44 ms, SE = 8.69 ms, t(298) = −5.10, p < 0.0001, 95% CI [−61 ms, −27 ms]), the linear mixed model revealed no significant effects of stimulation (β = 19 ms, SE = 0.046, t(298) = 0.42, p = 0.68, 95% CI [−71 ms, 109 ms]) or montage (β = −12 ms, SE = 0.026, t(298) = −0.45, p = 0.66, 95% CI [−63 ms, 40 ms]). Additionally, the lack of a significant two-way interaction effect between stimulation and montage (β = 15 ms, SE = 0.035, t(298) = 0.42, p = 0.68, 95% CI [−54 ms, 83 ms]) and a three-way interaction effect between time, stimulation, and montage (β = 6.85 ms, SE = 16.1 ms, t(298) = 0.43, p = 0.67, 95% CI [−25 ms, 39 ms]) indicated that neither the individual nor the combined effects of stimulation and montage influenced motor sequence learning trajectories over time (Figure 2).

In addition, the model indicated significant variability of the intercepts (main effect of intercept: β = 622 ms, SE = 34 ms, t(298) = 18.13, p < 0.0001, 95% CI [554 ms, 689 ms]) across individual subjects. This was successfully accounted for by including a random intercept for subjects (SD = 138 ms, 95% CI [113 ms, 168 ms]) in the linear mixed model. Also, the induced variability attributed to which sequence was practiced first, was negligible, as indicated by the random intercept for sequence (SD = 2.34e-8 ms). This confirmed that the specific sequence used had little to no impact on the outcome. Detailed model estimates and statistics are reported in the Supplementary Materials.

### Online improvements during stimulation

In addition to significant improvements over time when measured immediately after practice, participants also demonstrated online learning with concurrent stimulation (Figure 3), as evidenced by the significant main effect of time (β = −5.51 ms, SE = 0.396 ms, t(2269) = −13.92, p < 0.0001, 95% CI [−6.28 ms, −4.73 ms]) in the linear mixed model that included all sequenced blocks across the two practice sessions. Consistent with our previous findings, this linear mixed model also showed no overall significant effect of stimulation (main effect of stimulation: β = −18.53 ms, SE = 40.22 ms, t(2269) = −0.46, p = 0.65, 95% CI [−97.41 ms, 60.35 ms]) on online motor sequence learning. Interestingly, however, the model indicated an overall significant effect of montage (main effect of montage: β = −29.50 ms, SE = 7.41 ms, t(2269) = −3.98, p < 0.0001, 95% CI [−44.03 ms, −14.96 ms]) suggesting that, in general, subjects performed significantly better during practice sessions with the HD montage compared to sessions with the conventional montage, but independently of whether actual stimulation was delivered or not. In addition, the significant two-way interaction effect between time and montage (β = 2.61 ms, SE = 0.56 ms, t(2269) = 4.70, p < 0.0001, 95% CI [1.52 ms, 3.70 ms]) indicated that the rate of improvement over time also depended on montage, demonstrating that subjects showed steeper learning trajectories in sessions with the HD montage compared to sessions with the conventional montage. Moreover, the significant two-way interaction between stimulation and montage (β = 52.02 ms, SE = 9.88 ms, t(2269) = 5.27, p < 0.0001, 95% CI [32.65 ms, 71.39 ms]) revealed that the effect of stimulation was contingent on electrode montage, with stimulation during these sessions actually worsening performance.

**Figure 3.**
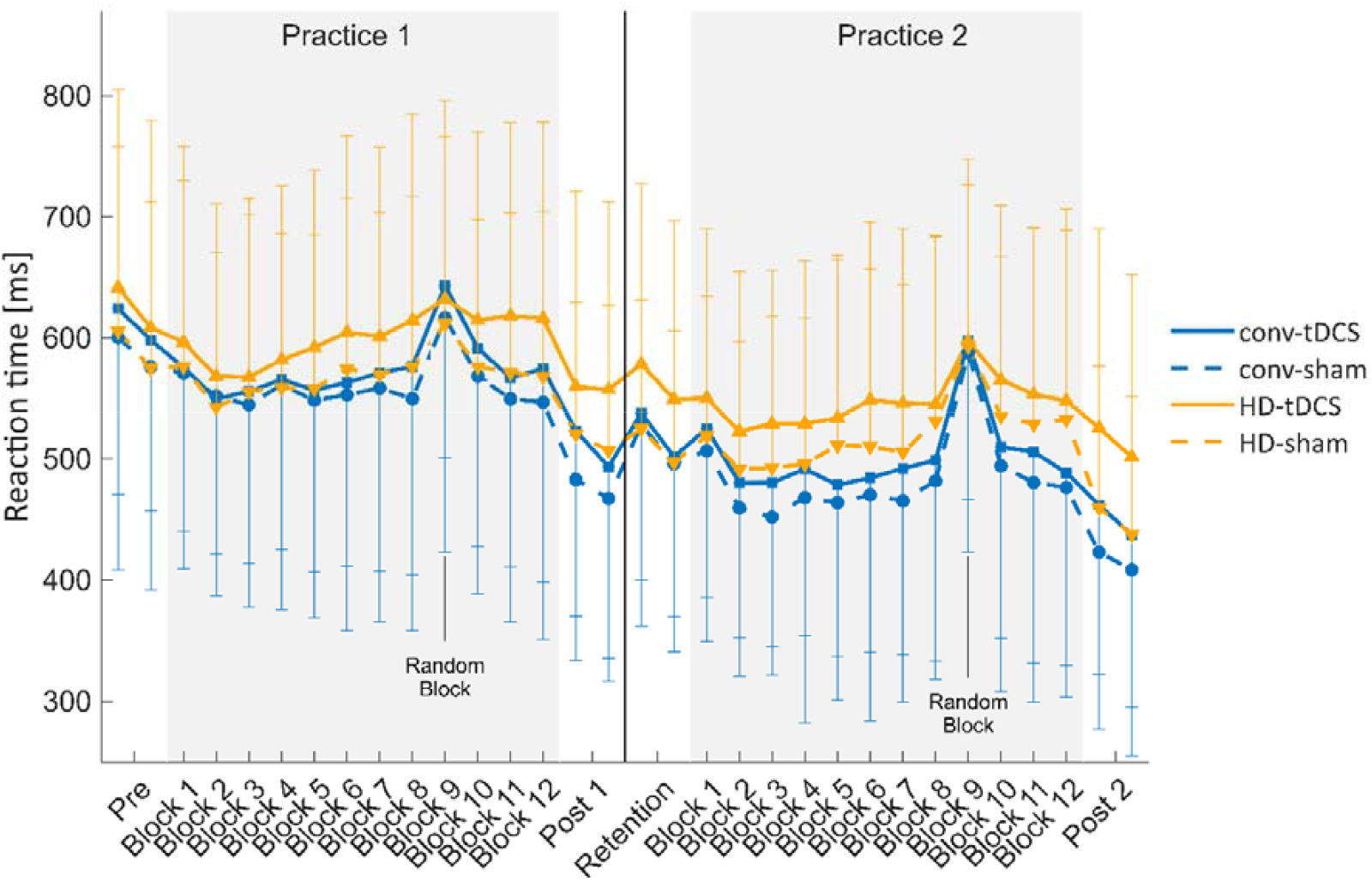
Online motor sequence learning. Mean reaction times in milliseconds [ms] during the serial reaction time task (SRTT) for) for conventional tDCS (dark blue), conventional sham (light blue), HD tDCS (orange), and HD sham (yellow). Data are presented as group means, and the error bars indicate the standard deviations.

Despite appearing contradictory at first, overall, these results demonstrated that conventional tDCS showed an apparent improvement in motor sequence learning by lowering the mean reaction times with on average −18.53 ms during online practice, though this was not statistically significant (main effect of stimulation: β = −18.53 ms, SE = 40.22 ms, t(2269) = −0.46, p = 0.65, 95% CI [−97.41 ms, 60.35 ms]). In contrast, whereas subjects seemed to perform generally better during practice sessions with the HD montage, independent of whether stimulation was applied (main effect of montage: β = −29.50 ms, SE = 7.41 ms, t(2269) = −3.98, p < 0.0001, 95% CI [−44.03 ms, −14.96 ms]), tDCS stimulation during these sessions appeared to worsen performance by increasing the mean reaction times by on average 33.5 ms (= −18.53 + 52.02) relative to sham. However, as we did not observe a triple interaction effect between stimulation, montage and time (β = −0.72 ms, SE = 0.74 ms, t(2269) = −0.97, p = 0.33, 95% CI [−2.18 ms, 0.73 ms]), the above mentioned effects did not change over time, meaning that nor conventional, nor HD tDCS significantly effected motor sequence learning as such.

Finally, and consistent with the random intercept effects reported in the previous section, the linear mixed model for online learning also demonstrated significant variability in the intercepts (main effect of intercept: β = 602 ms, SE = 36 ms, t(2269) = 16.80, p < 0.0001, 95% CI [532 ms, 672 ms]) across individual subjects. This was again successfully accounted for by including a random intercept for subjects (SD = 142 ms, 95% CI [117 ms, 172 ms]) in the model. In addition, the variability attributed to the sequence was minimal, as reflected in the small random intercept for SRTT sequence (SD = 27.6 ms, 95% CI [9.6 ms, 79.2 ms]), implying that the specific sequence had minimal impact. Detailed model estimates and statistics are reported in the Supplementary Materials.

### Continued learning in second practice session

Consistent with the first practice session, participants exhibited continued motor sequence learning in the second (Figure 2), as evidenced by the significant effect of time (main effect of time: β = −96 ms, SE = 14 ms, t(192) = −6.88, p < 0.0001, 95% CI [−124 ms, −69 ms]) when comparing performance at the end (Post 2) to the beginning of the second session (Retention). In line, the current linear mixed model likewise revealed no significant effect of stimulation (main effect of stimulation: β = −76 ms, SE = 75 ms, t(192) = −1.01, p = 0.312, 95% CI [−224 ms, 72 ms]) or montage (main effect of montage: β = −94 ms, SE = 69 ms, t(192) = −1.36, p = 0.176, 95% CI [−231 ms, 43 ms]), nor any two-way or three-way interaction effects between stimulation and time (β = 26 ms, SE = 19 ms, t(192) = 1.41, p = 0.159, 95% CI [−10 ms, 63 ms]), montage and time (β = 34 ms, SE = 20 ms, t(192) = 1.74, p = 0.084, 95% CI [−5 ms, 73 ms]), or stimulation montage and time (β = −14 ms, SE = 26ms, t(192) = −0.55, p = 0.585, 95% CI [−65 ms, 37 ms]).

In the second session, the inter-subject variability of performance at the beginning of the session (main effect of intercept: β = 804 ms, SE = 57 ms, t(192) = 14.23, p < 0.0001, 95% CI [693 ms, 915 ms]) was slightly larger than the inter-subject variability at the baseline in the beginning of the first session (main effect of intercept: β = 622 ms, SE = 34 ms, t(298) = 18.13, p < 0.0001, 95% CI [554 ms, 689 ms]). As before, we accounted for this variability by including the intercept for subjects as a random factor (random intercept for subjects: SD = 130 ms, 95% CI [106 ms, 158 ms]). Variability attributed to the different sequences that were practiced, was negligible (random intercept for SRTT sequences: SD = 8.08e-11 ms). The residual standard deviation was estimated at 0.045 ms (95% CI [0.041 ms, 0.051 ms]). Detailed model estimates and statistics are reported in the Supplementary Materials.

### Carryover effect

When assessing potential carryover effects from the crossover design, we found no evidence of a carryover effect as indicated by the nonsignificant effect of trial (main effect of trial: β = −116 ms, SE = 65 ms, t(95) = −1.78, p = 0.078, 95% CI [−245 ms, 13 ms]). Here, trial represented the blocks before (Trial 1: mean reaction time 632 ± 175 ms) and after crossover (Trial 2: mean reaction time 580 ± 139 ms). This means that prior exposure to stimulation or motor sequence learning in the first phase did not influence baseline performance in the second. Moreover, the model revealed no significant effects of stimulation (main effect of stimulation: β = −35 ms, SE = 61 ms, t(95) = −0.57, p = 0.573, 95% CI [−156 ms, 87 ms]) or montage (main effect of montage: β = −86 ms, SE = 65 ms, t(95) = −1.32, p = 0.191, 95% CI [-215 ms, 43 ms]), nor any significant interaction effects. The model did, however, reveal significant differences in the intercepts (main effect of intercept: β: 663 ms, SE = 47 ms, t(95) = 14.066, p < 0.0001, 95% CI [570 ms, 757 ms]. This variability was again accounted for by including the random intercept for subjects in the model (random intercept for subjects: SD = 147 ms, 95% CI [120 ms, 180 ms]). Viability attributable to the different sequences practiced was negligible (SD = 5.74e−6 ms). The residual standard deviation was only 49 ms, 95% CI [40 ms, 59 ms]) (see Supplementary Materials).

### Blinding

Subjects in the tDCS group reported significantly higher stimulation intensities on the VAS compared to the sham group (main effect of stimulation: p = 0.0080) and the reported intensity increased over time (main effect of time: p = 0.036) across the session. No significant effect of montage was observed (main effect of montage: p < 0.05). Most subjects in the tDCS group (70%) correctly identified that they received active tDCS throughout the experiment, while 20% believed they received no stimulation, and 10% reported uncertainty about the type of stimulation (Table 1). In the sham group, only 36% of the subjects correctly allocated themselves, while most (55%) believed to belong to the tDCS group. Only 1 subject indicated to be uncertain about the perceived type of stimulation. The experimenter correctly identified 70% of the tDCS subjects and 73% of the sham subjects, while only 23% were falsely allocated to the tDCS group.

**Table 1.** Results on the stimulation blinding assessment.

|  |  | Self-reported stimulation condition |  |  | Total |
| --- | --- | --- | --- | --- | --- |
| Stimulation condition |  | tDCS | sham | abstention |  |
| Subjects | tDCS | 21 | 6 | 3 | 30 |
|  | sham | 12 | 8 | 1 | 22 |
| Experimenter | tDCS | 21 | 9 | 0 | 30 |
|  | sham | 5 | 16 | 0 | 22 |

## Discussion

In this study, we aimed to investigate the effect of 1mA conventional anodal tDCS over M1 on motor sequence learning on retention 24h after one practice session compared to sham in healthy older adults. In addition, we evaluated the possible different effects of stimulation montage, i.e. HD tDCS compared to conventional tDCS, in a crossover design. We hypothesized that conventional anodal tDCS over M1 during motor sequence learning would be better able to improve and maintain performance after one practice session compared to sham stimulation in older adults. We based this expectation on our earlier work in both healthy older people and persons with Parkinson’s disease, even if these results were not always consistently positive.

Our findings showed that older adults exhibited improvements in motor sequence learning on the SRTT both within and across sessions, confirming the validity of our learning paradigm. However, the observed improvements were not significantly influenced by tDCS stimulation, regardless of whether a conventional or high-definition (HD) montage was used. While all groups showed improvements immediately after the first practice session and these gains were maintained after 24 hours, no additional consolidation effects were observed. Yet, all groups demonstrated continued learning during the second session, regardless of stimulation condition (active or sham) or montage type (conventional or HD).

Our results are consistent with some previous findings showing no effect of tDCS on motor sequence learning and retention, such as the results of Greeley et al. who were not able to show a significant effect after two practice sessions (Greeley et al. 2022). In line, King et al. 2020 found no significant neurophysiological changes at the group level following a single session of tDCS preceded by motor sequence learning in older adults (King et al. 2020). Interestingly, however, Ho et al. 2016 demonstrated that repeated sessions of tDCS led to a cumulative increase in cortical excitability (Ho et al. 2016), suggesting that sustained stimulation over multiple days induced measurable neuroplastic changes and behavioral effects. This suggests that multiple tDCS sessions may be necessary to enhance M1 excitability and induce lasting effects on motor sequence learning, with Dumel et al (2018) reporting SRTT performance improvements that persisted for up to three months following five sessions (Dumel et al. 2018). In addition, research in young adults demonstrated reliable effects on retention after multiple sessions of tDCS (Reis et al. 2015; Saucedo-Marquez et al. 2013; Waters-Metenier et al. 2014). This implies that one or two practice sessions may be insufficient to produce measurable improvements, which could account for the present results, despite evidence for single-session effects in prior study (Reis et al. 2009).

There are several other explanations for why we did not observe a significant effect of tDCS in our study. This may also relate to which extent M1 is engaged during motor sequence learning. Previous research indicates that the involvement of brain areas changes over time (Dahms et al. 2020), with M1 typically becoming increasingly important after multiple practice sessions. Consequently, tDCS targeting brain regions more strongly engaged during initial stages of motor sequence learning may have been more effective. Alternatively, applying tDCS over M1 at a later stage, when M1 is more strongly involved, may have been more likely to induce significant effects.

Another potential explanation may be related to the applied stimulation intensity and current density. Although unlikely, the stimulation intensity of 1 mA could have been insufficient to effectively modulate excitability in M1. A previous study demonstrated enhanced LTP-like plasticity following 3 mA of tDCS and when compared to lower intensities in older adults (Farnad et al. 2021). However, Ghasemian-Shirvan et al. 2023 did not find significant differences between 1, 2, and 3 mA during motor sequence learning on neither acquisition nor consolidation outcomes in a large sample of elderly. In fact, no significant effect of tDCS were observed, regardless of stimulation intensity (Ghasemian-Shirvan et al. 2023).

Interestingly, we found in an earlier power-based randomized double-blind sham-controlled study, which investigated the effects of boosting one session of writing training with tDCS of 1 mA in people with PD, better 24-hr retention and continued learning after tDCS and not after sham. This somewhat counterintuitive result may be explained by the specific neural profiles of people with PD, who typically display reduced motor learning consolidation (Cristini et al. 2023), as dopamine depletion leads to abnormal motor circuit activity and impaired synaptic plasticity (Sciacca et al. 2023). As such, tDCS may have helped to normalize or augment plasticity in the primary cortex, thereby boosting writing performance in PD. In contrast, older adults, who have a relatively intact dopaminergic systems, tDCS may not have added much or may have even disrupted a relatively balanced system. Another reason for the discrepant findings may be attributable to task-related differences, as recent meta-analysis in older adults with and without PD, demonstrated that tDCS effects are highly variable and task dependent (Vandendoorent et al. 2023). Whereas the previous study examined a writing task, the present study focused on motor sequence learning. Writing is particularly affected in PD, with amplitude deficits known as micrographia (Inzelberg et al. 2016), and may therefore have provided greater potential for improvement through tDCS-boosted motor learning, compared with motor sequence learning in healthy older adults.

It is also possible that the effectiveness of tDCS in older adults may have been impacted by electrophysiological changes in various brain regions in the cortex associated with aging (Habich et al. 2020). Older individuals exhibit, for example, reduced responsiveness to gamma-aminobutyric acid (GABA), a primary inhibitory neurotransmitter in the central nervous system that plays a key role in regulating excitability by maintaining the balance between excitation and inhibition of neurons. Considering that cortical excitability plays a major role in LTP-like plasticity and learning, changes in responsiveness to GABA can influence motor sequence learning in older adults. This is supported by Stagg et al. 2011, who showed that the responsiveness of GABA is correlated with motor learning capacity in healthy individuals (Stagg, Bachtiar, and Johansen-Berg 2011). In addition, they suggested that tDCS-induced changes in GABA levels may underly the effects of tDCS on motor sequence learning, as previous study has shown (Kim et al. 2014). As a result of the overall reduced responsiveness to GABA in older adults, tDCS may have elicited a differential neural response and was therefore less effective.

Anatomical alterations associated with aging could also affect tDCS effectiveness, as compared to young people. Evidence suggests that the thickness of the skull and cerebrospinal fluid change with age (Van Hoornweder et al. 2023; Yamada et al. 2023), and these changes could have a major impact on the magnitude of the resulting electric field at the cortical level (Laakso et al. 2015; Opitz et al. 2015). Whether these alterations significantly influence the effectiveness of tDCS depends on the espoused underlying neurophysiological mechanism, which still remains highly debated within the field (van Boekholdt et al. 2021). Assuming that tDCS exerts its effects through direct polarization of the membrane potentials of cortical neurons, referred to as the transcranial mechanism, skull thickness can have a significant impact on stimulation efficacy. As the tDCS current passes through the scalp, skull, and cerebrospinal fluid, the resulting cortical electric field is attenuated accordingly. Consequently, a thicker skull may lead to greater attenuation of the electric field. Alternatively, if tDCS exerts its effects primarily through indirect mechanisms, as suggested by our group (van Boekholdt et al. 2021), factors like skull thickness may have less impact on stimulation efficacy.

Under the assumption that tDCS elicits its behavioral effects by increasing arousal and vigilance through peripheral mechanisms (van Boekholdt et al. 2021), electrode type and montage may not exert as much influence on behavioral outcomes as traditionally presumed. Many researchers have previously attributed the limited beneficial effects of conventional tDCS to suboptimal targeting of the relevant brain regions (Schmidt et al. 2024). However, although many studies indicated that the HD montage elicits more probing effects compared to the conventional tDCS montage, recent findings challenge the idea that greater anatomical precision necessarily leads to better results. For instance, a recent study found that conventional M1-tDCS improved sequence learning, whereas HD tDCS impaired performance (Firouzi et al. 2023b). Our short-term learning effects partly confirm these findings. The authors also suggested that HD tDCS may fail to sufficiently engage broader neural networks involved in sequence learning, in contrast to the more diffuse stimulation provided by conventional tDCS. Similarly, Kaminski et al. 2024 found that HD tDCS over DLPFC or M1 did not enhance performance in a complex motor task (Kaminski et al. 2024), suggesting that focal stimulation may not reach the distributed networks involved. An alternative explanation for the lack of observed effects in this study, even showing some detriment of HD on performance, could be the inhibitory effect on the return electrodes in the HD montage, which may have induced unintended suppressive effects. Together, these studies highlight that broader stimulation from conventional montages may be more effective than focal HD tDCS in modulating tasks requiring broader neural networks. However, all the above explanations for the negligible effects of stimulation in our study remain speculative.

Although this study was preregistered and we performed a priori power analysis, it is important to mention a few limitations. First, despite using permuted block randomization for group assignment, unevenly distributed dropouts after inclusion resulted in unequal stimulation group sizes. While this imbalance may not be ideal, they are unlikely to have influenced our findings. Second, most subjects in the tDCS group were able to correctly identify that they had received active tDCS and therefore despite our efforts to have blinded procedures, blinding was not fully successful. However, since no significant differences were observed between groups, it is unlikely that this affected the outcomes of our study. Finally, although no group differences were observed in demographic measures, the study included subjects across a relatively wide age range and this variability could have introduced individual differences in learning performance. Future studies may benefit from more homogeneous samples to better isolate and understand potential age-related effects on learning and stimulation effects.

In summary, while earlier studies reported beneficial effects of tDCS (Reis et al. 2015; Saucedo-Marquez et al. 2013; Waters-Metenier et al. 2014), even with a single session (Reis et al. 2009), our findings indicated that the potential effects of tDCS on motor sequence learning in older adults, are more limited and variable than the literature suggests, regardless of whether a conventional or high-definition (HD) montage was used.

## Conclusion

Our results indicated that neither conventional tDCS nor HD tDCS was able to significantly enhance motor sequence learning in older adults compared to sham. These results align with a broader set of studies in the tDCS literature questioning the replicability of earlier claims regarding tDCS effects on motor learning and other behavioral outcomes. This highlights the need for a better understanding of the neurophysiological mechanisms of tDCS, and how individual and aging-related neurophysiological differences affect responsiveness. Future research should explore whether multiple sessions of tDCS can elicit a more reliable effect as well as clarify if and under which conditions tDCS can be used as a tool for noninvasive neuromodulation to supplement motor learning.

## Acknowledgment

Internal Funds of the KU Leuven (C14/17/115 to AN and JJOdX) supported this study; SK and SB were FWO doctoral researchers (resp. FWO grant numbers S32421N and 1167419N). All funders had no role in study design, data collection and analysis, decision to publish, or preparation of the manuscript.

## Author contribution

1. Research project: A. Conception, B. Organization, C. Execution; 2. Data: A. Acquisition, B. Processing; 3. Statistical Analysis: A. Design, B. Execution, C. Review and Critique; 4. Manuscript: A. Writing, B. Review

SK: 2B, 3A, 3B, 3C, 4A, 4B

BV: 1A, 1B, 1C, 2A, 2B, 3A, 3B, 4A

SB: 1A, 1B, 1C, 2A, 2B, 3A, 3B, 3C, 4B

MG: 1A, 3A, 4B

EN: 1A, 1B, 1C, 3A, 3C, 4B

JO: 1A, 1B, 1C, 3A, 3C, 4B

AN: 1A, 1B, 1C, 3A, 3C, 4B

## Acknowledgment

We thank all the people who participated in our study.

## Supplementary materials

### Subjects

Fifty-two subjects were randomized to either the tDCS (n = 30) group or the sham (n = 22) group (Table S1). All subjects were cognitively intact, as indicated by Montreal Cognitive Assessment (MoCA) scores averaging above 26 in both groups. Statistical comparisons indicated no group differences in demographic or neuropsychological measures, including age, gender, cognition (MoCA), executive function and processing speed (TMT-A and TMR-B), sleep quality (PSQI), mood (HADS Anxiety and HADS Depression), and manual dexterity (DEXTQ-24), with all p > 0.2.

**Table S1.**
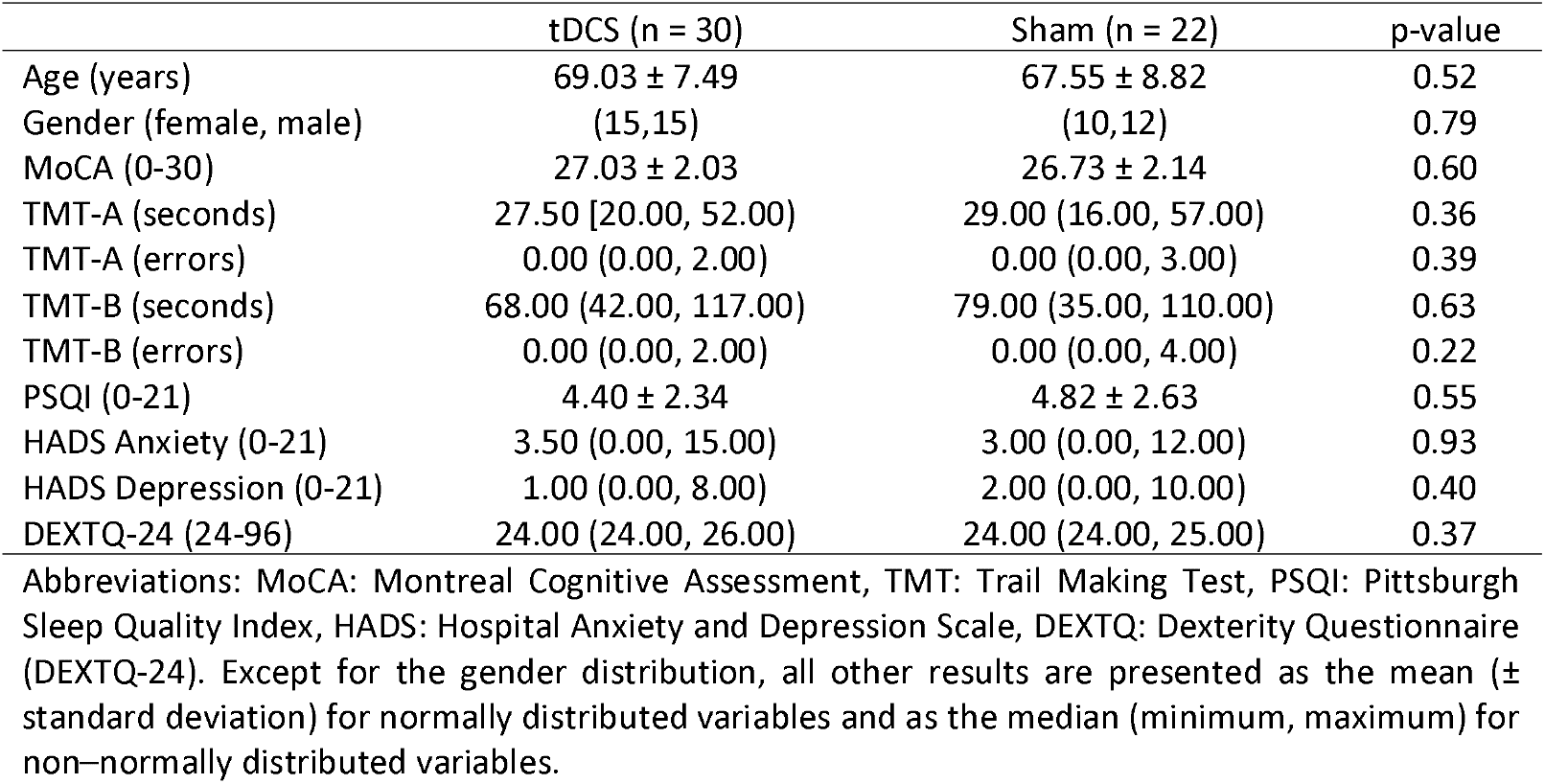
Subject characteristics.

|  | tDCS (n = 30) | Sham (n = 22) | p-value |
| --- | --- | --- | --- |
| Age (years) | 69.03 ± 7.49 | 67.55 ± 8.82 | 0.52 |
| Gender (female, male) | (15,15) | (10,12) | 0.79 |
| MoCA (0-30) | 27.03 ± 2.03 | 26.73 ± 2.14 | 0.60 |
| TMT-A (seconds) | 27.50 [20.00, 52.00] | 29.00 (16.00, 57.00) | 0.36 |
| TMT-A (errors) | 0.00 (0.00, 2.00) | 0.00 (0.00, 3.00) | 0.39 |
| TMT-B (seconds) | 68.00 (42.00, 117.00) | 79.00 (35.00, 110.00) | 0.63 |
| TMT-B (errors) | 0.00 (0.00, 2.00) | 0.00 (0.00, 4.00) | 0.22 |
| PSQI (0-21) | 4.40 ± 2.34 | 4.82 ± 2.63 | 0.55 |
| HADS Anxiety (0-21) | 3.50 (0.00, 15.00) | 3.00 (0.00, 12.00) | 0.93 |
| HADS Depression (0-21) | 1.00 (0.00, 8.00) | 2.00 (0.00, 10.00) | 0.40 |
| DEXTQ-24 (24-96) | 24.00 (24.00, 26.00) | 24.00 (24.00, 25.00) | 0.37 |

Abbreviations: MoCA: Montreal Cognitive Assessment, TMT: Trail Making Test, PSQI: Pittsburgh Sleep Quality Index, HADS: Hospital Anxiety and Depression Scale, DEXTQ: Dexterity Questionnaire (DEXTQ-24). Except for the gender distribution, all other results are presented as the mean (± standard deviation) for normally distributed variables and as the median (minimum, maximum) for non-normally distributed variables.

### Linear mixed model outcomes

#### Effect of stimulation and montage on learning and retention

The linear mixed model is used to test the effects of stimulation and montage on motor sequence learning and retention over time as reflected in serial reaction time task (SRTT) mean reaction times (RT) per block across the following three time points: baseline performance (Pre), performance immediately after the first practice session (Post 1) and performance 24h after practice (Retention).

Outcome Variable:

Mean reaction time (RT) per block in the serial reaction time task (SRTT) per block

Fixed Effects:

Stimulation (active tDCS vs. sham stimulation)
Montage (conventionnel vs. HD montage)

Time: Blocks

Pre (Block 1)
Post (Block 2)
Retention (Block 3)

Random Effects:

Subject
SRTT sequence

Linear mixed-effects model fit by maximum likelihood:

Formula:

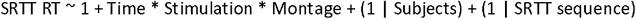

Model information:

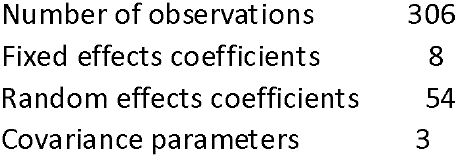

Model fit statistics:

| AIC | BIC | LogLikelihood | Deviance |
| --- | --- | --- | --- |
| -681.99 | -641.04 | 352 | -703.99 |

Fixed effects coefficients (95% CIs):

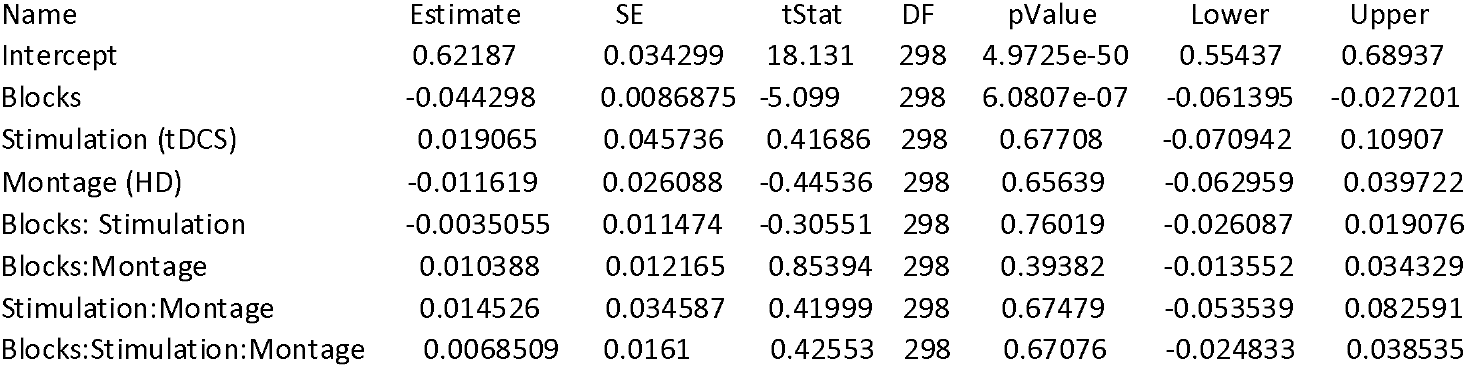

| Name | Estimate | SE | tStat | DF | pValue | Lower | Upper |
| --- | --- | --- | --- | --- | --- | --- | --- |
| Intercept | 0.62187 | 0.034299 | 18.131 | 298 | 4.9725e-50 | 0.55437 | 0.68937 |
| Blocks | -0.044298 | 0.0086875 | -5.099 | 298 | 6.0807e-07 | -0.061395 | -0.027201 |
| Stimulation (tDCS) | 0.019065 | 0.045736 | 0.41686 | 298 | 0.67708 | -0.070942 | 0.10907 |
| Montage (HD) | -0.011619 | 0.026088 | -0.44536 | 298 | 0.65639 | -0.062959 | 0.039722 |
| Blocks: Stimulation | -0.0035055 | 0.011474 | -0.30551 | 298 | 0.76019 | -0.026087 | 0.019076 |
| Blocks:Montage | 0.010388 | 0.012165 | 0.85394 | 298 | 0.39382 | -0.013552 | 0.034329 |
| Stimulation:Montage | 0.014526 | 0.034587 | 0.41999 | 298 | 0.67479 | -0.053539 | 0.082591 |
| Blocks:Stimulation:Montage | 0.0068509 | 0.0161 | 0.42553 | 298 | 0.67076 | -0.024833 | 0.038535 |

Random effects covariance parameters (95% CIs): Group: Subjects (52 Levels)

| Name1 | Name2 | Type | Estimate | Lower | Upper |
| --- | --- | --- | --- | --- | --- |
| Intercept | Intercept | std | 0.13798 | 0.1132 | 0.1682 |

Group: SRTT sequences (2 Levels)
| Name1 | Name2 | Type | Estimate | Lower | Upper |
| --- | --- | --- | --- | --- | --- |
| Intercept | Intercept | std | 2.3441e-11 | NaN | NaN |

Group: Error
| Name | Estimate | Lower | Upper |
| --- | --- | --- | --- |
| Residual Std | 0.056493 | 0.051787 | 0.061627 |

#### Continued learning in second practice session

The linear mixed model was used to test if subjects showed continued learning in the second practice session and to investigate the potential effects of stimulation and montage on continued motor sequence learning as reflected by mean reaction times (RT) per block in the serial reaction time task (SRTT) across the following two time points: at the beginning of the second session (Retention), and at the end of the second practice session (Post 2).

Outcome Variable:

Mean reaction time (RT) per block in the serial reaction time task (SRTT)

Fixed Effects:

Stimulation (active tDCS vs. sham stimulation)
Montage (conventional vs. HD montage)

Random Effects:

Subject
SRTT sequence

Time: Blocks

Retention (Block 3)
Post 2 (Block 4)

Linear mixed-effects model fit by maximum likelihood: Formula:

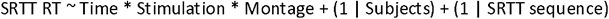

Model information:

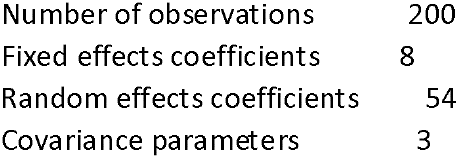

Model fit statistics:

| AIC | BIC | LogLikelihood | Deviance |
| --- | --- | --- | --- |
| -466.95 | -430.67 | 244.48 | -488.95 |

Fixed effects coefficients (95% CIs):

| Name | Estimate | SE | tStat | DF | pValue | Lower | Upper |
| --- | --- | --- | --- | --- | --- | --- | --- |
| Intercept | 0.804 | 0.056502 | 14.23 | 192 | 7.5994e-32 | 0.69256 | 0.91545 |
| Blocks | -0.096406 | 0.014021 | -6.8759 | 192 | 8.4226e-11 | -0.12406 | -0.068751 |
| Stimulation (tDCS) | -0.076007 | 0.074943 | -1.0142 | 192 | 0.31177 | -0.22382 | 0.071811 |
| Montage (HD) | -0.094192 | 0.069351 | -1.3582 | 192 | 0.176 | -0.23098 | 0.042597 |
| Blocks: Stimulation | 0.026239 | 0.018548 | 1.4147 | 192 | 0.15878 | -0.010344 | 0.062823 |
| Blocks: Montage | 0.034014 | 0.019602 | 1.7353 | 192 | 0.0843 | -0.0046483 | 0.072677 |
| Stimulation: Montage | 0.081328 | 0.091832 | 0.88561 | 192 | 0.37694 | -0.099802 | 0.26246 |
| Blocks: Stimulation: Montage | -0.014185 | 0.025962 | -0.54639 | 192 | 0.58543 | -0.065392 | 0.037022 |

Random effects covariance parameters (95% CIs):

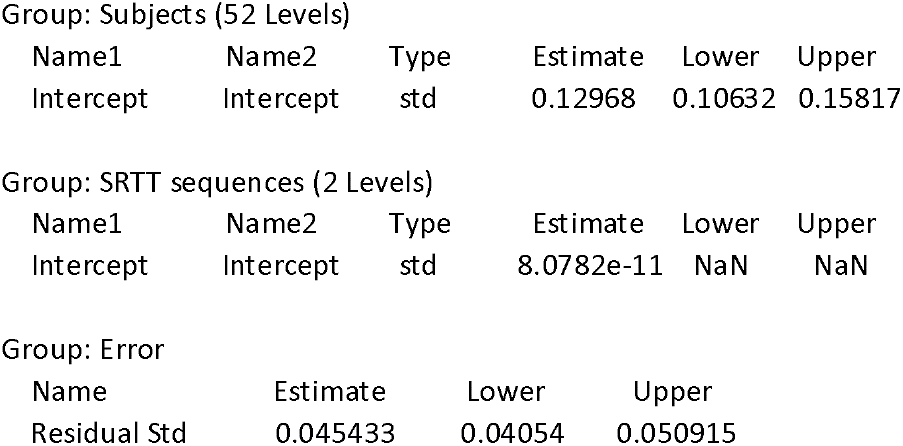

Random effects covariance parameters (95% CIs): Group: Subjects (52 Levels)
| Name1 | Name2 | Type | Estimate | Lower | Upper |
| --- | --- | --- | --- | --- | --- |
| Intercept | Intercept | std | 0.12968 | 0.10632 | 0.15817 |

Group: SRTT sequences (2 Levels)
| Name1 | Name2 | Type | Estimate | Lower | Upper |
| --- | --- | --- | --- | --- | --- |
| Intercept | Intercept | std | 8.0782e-11 | NaN | NaN |

Group: Error
| Name | Estimate | Lower | Upper |
| --- | --- | --- | --- |
| Residual Std | 0.045433 | 0.04054 | 0.050915 |

#### Online improvements during stimulation

The linear mixed model was used to investigate the effects of stimulation and montage on online motor sequence learning during learning and concurrent stimulation as reflected by the mean reaction times (RT) per block of the serial reaction time task (SRTT) including all sequenced blocks across the two learning sessions, while excluding the random sequence block (Block 9) to avoid its influence on the online learning trajectories.

Outcome Variable:

Mean reaction time (RT) per block in the serial reaction time task (SRTT)

Fixed Effects:

Stimulation (active tDCS vs. sham stimulation)
Montage (conventional vs. HD montage)

Time: Blocks

All sequenced blocks across the two practice sessions

Random Effects:

Subject
SRTT sequence

Linear mixed-effects model fit by maximum likelihood: Formula:

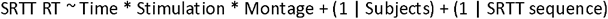

Model information:

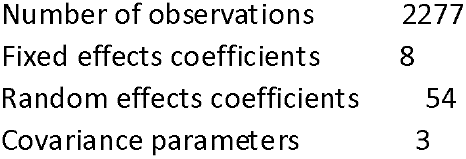

Model fit statistics:

| AIC | BIC | LogLikelihood | Deviance |
| --- | --- | --- | --- |
| -6368.2 | -6305.2 | 3195.1 | -6390.2 |

Fixed effects coefficients (95% CIs):

| Name | Estimate | SE | tStat | DF | pValue | Lower | Upper |
| --- | --- | --- | --- | --- | --- | --- | --- |
| Intercept | 0.60208 | 0.035835 | 16.801 | 2269 | 8.4357e-60 | 0.53181 | 0.67235 |
| Blocks | -0.0055058 | 0.00039569 | -13.915 | 2269 | 2.6866e-42 | -0.0062818 | -0.0047299 |
| Stimulation (tDCS) | -0.018528 | 0.040223 | -0.46063 | 2269 | 0.64511 | -0.097405 | 0.060349 |
| Montage (HD) | -0.029496 | 0.0074133 | -3.9788 | 2269 | 7.1446e-05 | -0.044033 | -0.014958 |
| Blocks: Stimulation | 0.0007447 | 0.00052662 | 1.4141 | 2269 | 0.15747 | -0.00028801 | 0.0017774 |
| Blocks: Montage | 0.0026129 | 0.00055573 | 4.7017 | 2269 | 2.7346e-06 | 0.0015231 | 0.0037026 |
| Stimulation: Montage | 0.052017 | 0.009879 | 5.2655 | 2269 | 1.5299e-07 | 0.032645 | 0.07139 |
| Blocks: Stimulation: Montage | -0.00072276 | 0.00074185 | -0.97426 | 2269 | 0.33003 | -0.0021775 | 0.00073203 |

Random effects covariance parameters (95% CIs): Group: Subjects (52 Levels)

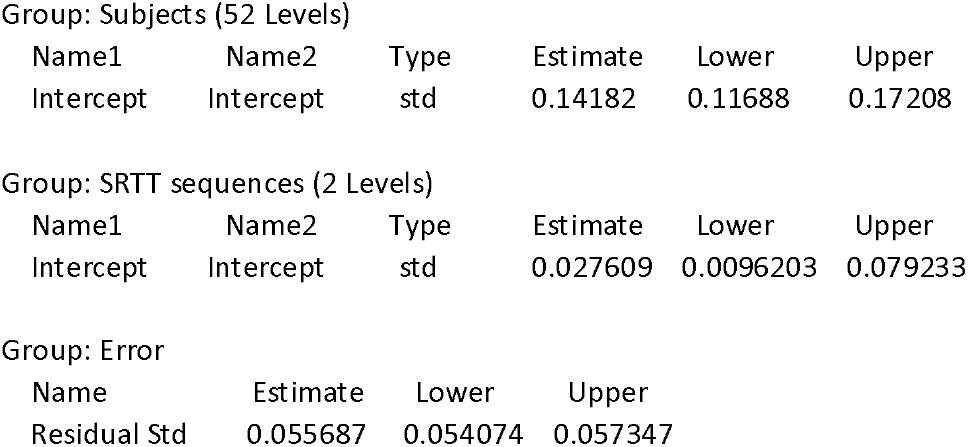

#### Carryover effect

A linear mixed model was used to investigate potential carryover effects resulting from the crossover design by comparing the mean reaction times (RTs) in the first two sequenced blocks (Pre) of the first sessions of the serial reaction time task (SRTT) before (Trial 1) and after (Trial 2) crossover.

Outcome Variable:

Mean reaction time (RT) per block in the serial reaction time task (SRTT)

Fixed Effects:

Stimulation (active tDCS vs. sham stimulation)
Montage (conventional vs. HD montage)

Random Effects:

Subject
SRTT sequence

Time: Trail

Pre before crossover (Trial 1)
Pre after crossover (Trial 2)

Linear mixed-effects model fit by maximum likelihood: Formula:

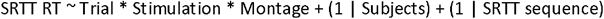

Model information:

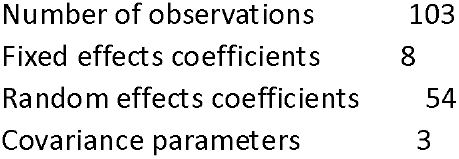

Model fit statistics:

| AIC | BIC | LogLikelihood | Deviance |
| --- | --- | --- | --- |
| -154.91 | -125.93 | 88.454 | -176.91 |

Fixed effects coefficients (95% CIs):

| Name | Estimate | SE | tStat | DF | pValue | Lower | Upper |
| --- | --- | --- | --- | --- | --- | --- | --- |
| Intercept | 0.66339 | 0.047161 | 14.066 | 95 | 5.894e-25 | 0.56976 | 0.75701 |
| Trial (Trial 2) | -0.11592 | 0.065003 | -1.7832 | 95 | 0.077741 | -0.24496 | 0.013131 |
| Stimulation (tDCS) | -0.034561 | 0.061033 | -0.56627 | 95 | 0.57254 | -0.15573 | 0.086605 |
| Montage (HD) | -0.085612 | 0.065003 | -1.3171 | 95 | 0.19099 | -0.21466 | 0.043434 |
| Trial:Stimulation | 0.077225 | 0.087025 | 0.88739 | 95 | 0.37711 | -0.095541 | 0.24999 |
| Trial:Montage | 0.14262 | 0.12629 | 1.1293 | 95 | 0.26161 | -0.10809 | 0.39332 |
| Stimulation:Montage | 0.13738 | 0.087025 | 1.5787 | 95 | 0.11774 | -0.035383 | 0.31015 |
| Trial:Stimulation:Montage | -0.20425 | 0.16934 | -1.2061 | 95 | 0.23076 | -0.54043 | 0.13194 |

Random effects covariance parameters (95% CIs): Group: Subjects (52 Levels)

| Name1 | Name2 | Type | Estimate | Lower | Upper |
| --- | --- | --- | --- | --- | --- |
| Intercept | Intercept | std | 0.14708 | 0.11996 | 0.18033 |

Group: SRTT sequences (2 Levels)
| Name1 | Name2 | Type | Estimate | Lower | Upper |
| --- | --- | --- | --- | --- | --- |
| Intercept | Intercept | std | 5.7371e-09 | NaN | NaN |

Group: Error
| Name | Estimate | Lower | Upper |
| --- | --- | --- | --- |
| Residual Std | 0.048796 | 0.040168 | 0.059276 |

